# Detecting environment-dependent diversification from phylogenies: a simulation study and some empirical illustrations

**DOI:** 10.1101/162248

**Authors:** Eric Lewitus, Hélène Morlon

## Abstract

Understanding the relative influence of various abiotic and biotic variables on diversification dynamics is a major goal of macroevolutionary studies. Recently, phylogenetic approaches have been developed that make it possible to estimate the role of various environmental variables on diversification using time-calibrated species trees, paleoenvironmental data, and maximum-likelihood techniques. These approaches have been effectively employed to estimate how speciation and extinction rates vary with key abiotic variables, such as temperature and sea level, and we can anticipate that they will be increasingly used in the future. Here we compile a series of biotic and abiotic paleodatasets that can be used as explanatory variables in these models and use simulations to assess the statistical properties of the approach when applied to these paleodatasets. We demonstrate that environment-dependent models perform well in recovering environment-dependent speciation and extinction parameters, as well as in correctly identifying the simulated environmental model when speciation is environment-dependent. We explore how the strength of the environment-dependency, tree size, missing taxa, and characteristics of the paleoenvironmental curves influence the performance of the models. Finally, using these models, we infer environment-dependent diversification in three empirical phylogenies: temperature-dependence in Cetacea, *δ*^13^*C*-dependence in Ruminantia, and *CO*_2_-dependence in Portulacaceae. We illustrate how to evaluate the relative importance of abiotic and biotic variables in these three clades and interpret these results in light of macroevolutionary hypotheses for mammals and plants. Given the important role paleoenvironments are presumed to have played in species evolution, our statistical assessment of how environment-dependent models behave is crucial for their utility in macroevolutionary analysis.

---

For decades, evolutionary biologists and paleontologists have debated the relative role of major biotic and abiotic drivers of macroevolutionary dynamics (76; (84). The view that links biodiversity to abiotic change, which has been apparent to biologists since the 19th century (83; (85) and is generally referred to as the Court Jester hypothesis (3), sees shifts in diversification as timed with climatic or geologic events. The broad-scale synchrony between global biodiversity and temperature throughout the Phanerozoic (72; (49) supports this view. More focused observations, such as the rise of angiosperms during the early Cretaceous thermal maximum (77; (18), also find support for close ties between abiotic change and species diversification. The view that links biodiversity shifts to biotic interactions, on the other hand, as outlined, for example, in Leigh Van Valen‘s Red Queen hypothesis (80) and Ehrlich and Raven‘s Escape and Radiate hypothesis (24), suggests that interspecific interactions (e.g., predation, mutualism, and species recognition) drive species diversification (47). While these various abiotic and biotic factors offer fundamentally different interpretations of how species evolve over geological time, evaluating their relative importance in practice is not trivial and has been inhibited by methodological shortcomings.

Investigations into the drivers of species diversification have been conducted using fossil specimens and phylogenetic comparative methods. Time-series of fossil specimens have been used to study speciation and extinction rates through time (70; (73), the timing and effect of mass extinction events on species richness (67;25;15;46), and correlations between environmental changes and shifts in biodiversity (8; 28; 35). Fossil studies remain key to our understanding of macroevolutionary dynamics, but they are hindered by incomplete sampling (75), clumping of specimens near mass extinction events (39), and high sensitivity to taxonomic and spatial scale (51; (26). Importantly, fossil studies are usually restricted to the few lineages for which sufficient fossil information is available (7). Phylogenetic comparative methods, which tender statistical or probabilistic information from species trees, provide an alternative approach to the same type of questions, applicable to a more diverse set of organisms. They have been used to study patterns of speciation and extinction in clades (55; (52), as well as how these patterns of diversification are related to paleoenvironmental drivers (78; 87; 20; 12; 42; 16; 21; 61).

Until recently, phylogenetic approaches for analyzing the role of paleoenvironmental drivers relied on correlative techniques (20; 78; 87), often assuming that environmental changes are punctuated events. This has been improved in the last couple of years, with the development of ‘environment-dependent’ likelihood-based approaches (19), implemented in the R package RPANDA (53), that allow testing whether gradual changes in paleoenvironments had a significant influence on speciation and extinction rates, as well as quantifying the direction and magnitude of this potential influence.

Environment-dependent models take the form of time-dependent birth-death models (54), but where the speciation and extinction rates, λ(*t*) and *μ*(*t*), respectively, at a given time, *t*, can depend on measured time-variable palaeodata, *E*(*t*). In order to test the influence of the paleoenvironment on diversification, hypotheses are made about the functional form of the dependency. Two main dependencies have been used: a linear dependency, of the form:

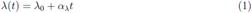

and

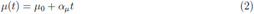

and an exponential dependency of the form

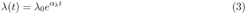

and

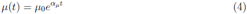

where λ_0_, *μ*_0_, *α*_λ_, and *α*_*μ*_ are free parameters. The probability density of observing a set of branching times in a phylogeny with a specific environmental dependence can then be computed using the likelihood expressions derived for time-dependent models, adapted to accommodate a time-dependent environmental variable (54; (19). The significance of the environmental dependence, its form (e.g. linear or exponential), its sign (positive or negative), and its strength can thus be assessed using maximum likelihood techniques.

Likelihood-based phylogenetic comparative methods have been essential aids to uncovering patterns of macroevolutionary change (52), and we can thus anticipate that the environment-dependent models will be of great use for understanding the relative role of major biotic and abiotic drivers on deep-time diversity dynamics. These models have already identified, for example, a positive relationship between speciation rates and global temperature in ruminants in the Cenozoic (12), an inverse relationship between net diversification rates and temperature in bird species since the KpG boundary (16), a positive relationship between extinction rates and sea level in birdwing butterflies (21), and a positive relationship between speciation rates and Andean uplift in neotropical orchids (61). However, we can only have a relative confidence in these empirical results, because the performances of the environment-dependent models have not been formally assessed. We can be confident that the likelihood expressions are correct, because the environment-dependent models are straightforward extensions of time-dependent models that have themselves been thoroughly tested using simulations (54; (31); but this does not guarantee that parameter estimation on reasonably sized phylogenies is not biased, that there is enough statistical power in the data to distinguish between alternative evolutionary scenarios (such as alternative environmental dependences), or that environmental dependence is not artifactually inferred when it did not occur. So, for example, temperature-dependency of birds diversification (16) was inferred on a highly undersampled phylogeny (*~* 0.02%); even though the likelihoods accommodate missing taxa, can such results be trusted? Environment-dependent extinction was detected in birds (16) and birdwing butterflies (21). Can we trust these results given the well-known difficulty in inferring extinction rates from reconstructed phylogenies (52)? We cannot answer these questions, or similar questions that will undoubtedly arise in future use of environment-dependent models, unless the statistical properties of these models are thoroughly assessed using simulations.

In this paper, we compile four biotic and five abiotic paleodatasets that can potentially be used as explanatory variables in environment-dependent models and use simulated trees to evaluate the statistical performance of the corresponding models. We first focus on temperature-dependent models: we test the ability to recover speciation rates, extinction rates, and their temperature-dependencies; we also test the ability to detect temperature-dependent diversification when it occurs and check that such dependency is not wrongly detected when it does not occur. Next, because a particularly promising use of these models is to find which environmental variable, among a set, best explains diversification dynamics, we test the ability to correctly identify the environmental variable that actually influenced diversification rates and assess whether specific characteristics of the paleoenvironmental curves influence this ability. Finally, we offer best-practice guidelines for using the environment-dependent model and implement them on empirical phylogenies.

## Materials and Methods

### Paleodata

We collected paleodata on nine variables that have been associated with macroevolutionary hypotheses of diversification. These paleodata represent only a subset of the paleoenvironments that could potentially be used in environment-dependent diversification models, and are focused primarily on the marine environment. However, most of them are also relevant to the terrestrial environment; and they cover a wide range of biotic and abiotic hypotheses with minimal redundancy. Most importantly, they cover a wide range of resolution and temporal trends; we expect the statistical properties of the environment-dependent models on these paleodata to provide a good representation of their statistical properties in general. The paleodata can roughly be classified into three main categories. (i) Paleoclimatic variables: temperature data (inferred from *δ*^18^*O* measurements) taken from (89); atmospheric *CO*_2_ data taken from (37); benthic *δ*^13^*C* data taken from (45). Temperature is the canonical indicator of climate change. Atmospheric *CO*_2_ varies as a result of biotic changes, such as the rise of photosynthetic plankton (60), or tectonic activity, which causes either a reduction in *CO*_2_ caused by the subduction of carbonate-rich ocean crust or the emission of *CO*_2_ through volcanic eruptions (5); levels of atmospheric *CO*_2_ are thought to impact, in particular, photosynthetic organisms. *δ*^13^*C* reflects global changes in organic carbon sequestration and organic isotope fractionation ratios. In the Cenozoic, *δ*^13^*C* can represent changes in the proportion of *C*_4_ and *C*_3_ grasses dominating the planet, and is therefore likely to be crucial to the evolution of grasses and the herbivores that feed on them; *δ*^13^*C* can also represent increases in ^13^*C*-rich marine species, such as diatoms, which contribute considerably to the overall carbon present in the planktonic food web (40; (59). (ii) Variables reflecting changes in oceanic composition: silica weathering ratio (14) and sea level estimates (50). The silica weathering ratio accounts for influxes of silicic acid in the ocean, which is needed for silica-precipitating microalgae (e.g., diatoms and radiolaria), which are responsible for about a quarter of global primary production (10). Sea level has several effects on both the terrestrial and the marine biome, through, for example, different overall land surfaces, different weathering levels, changes in stoichiometry, and currents. (iii) Variables reflecting interspecific interactions (primarily in the ocean): we constructed diversity curves for fossil archaeplastida, radiolaria, foraminifera, and ostracods. Fossil data were compiled from the Neptune Database (44) and Paleobiology Database (https://paleobiodb.org/) and diversity curves were estimated at the genus level using shareholder quorum subsampling (1) at two-million-year bins. These diversity curves are useful for testing hypotheses of direct species interactions, such as grazing of archaeplastida by herbivores, or indirect interactions, such as competition for the same feeding resources; they may also be useful, for example, as proxies for global productivity (archaeplastida, (68)). Additionally, these fossil species were chosen because they are abundant enough in the fossil record to generate detailed diversity curves.

All paleoenvironmental curves were stored in the R package RPANDA freely available on CRAN (53). For the purpose of our study, to avoid biases in model selection, curves were truncated at 52 million years ago (the earliest date of the silica curve) and values were scaled to between 0 and 1.

### Performance of temperature-dependent diversification models

We used simulations to test the statistical properties of the temperature-dependent diversification models. All our simulations were conducted over the time span of the environmental curves (i.e., 52 Myrs). We discarded trees with fewer than 50 tips, which always corresponded to less than 10% of the simulated trees. Throughout, speciation and extinction rates are in units of events per million years.

We tested the ability of temperature-dependent models to recover accurate parameter values. We simulated environment-dependent birth-death trees with speciation rates, λ, and/or extinction rates, *μ*, that varied as an exponential function of temperature T, such that at each given point in time, 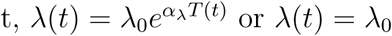 and 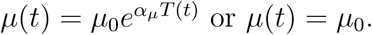 For trees with a λ-dependency on T (hereafter referred to as temperature-dependent λ trees), we simulated with λ_0_ = 0.1, 0.15, 0.2, fixing *α*_λ_ = 0.02 and *μ*_0_ = 0.01; *α*_λ_ = *-*0.5, -0.2, 0.4, 0.8, 1.6 fixing λ_0_ = 0.1 and *μ*_0_ = 0.01; and *μ*_0_ = 0.01, 0.03, 0.06, 0.09 fixing λ_0_ = 0.1 and *α*_λ_ = 0.02. We furthermore simulated temperature-dependent λ trees with a considerably higher *μ* (*μ*_0_ = 0.7) to reflect estimates of *μ* inferred from the fossil record (32). To do so, we increased λ too (λ_0_ = 0.7, *α*_λ_ = 0.35), otherwise it was difficult to simulate surviving trees. For trees with a *μ*-dependency on T (hereafter referred to as temperature-dependent *μ* trees), we simulated with λ_0_ = 0.2, 0.25, 0.3, fixing *μ*_0_ = 0.02 and *α*_*μ*_ = 0.02; *μ*_0_ = 0.02, 0.06, 0.1, fixing λ_0_ = 0.3 and *α*_*μ*_ = 0.01; and *α*_*μ*_ = *-*1, -0.6, 0.2, 0.4, fixing λ_0_ = 0.5 and *μ*_0_ = 0.1. Additionally, we simulated trees with both a λ- and *μ*-dependency on T, with (1) λ_0_ = 0.05,*α*_λ_ = *-*0.15, 0.15, *μ*_0_ = 0.05, and *α*_*μ*_ = 0.1; (2) λ_0_ = 0.25, *α*_λ_ = 0.4, *μ*_0_ = 0.05, and *α*_*μ*_ = 0.1; (3) λ_0_ = 0.02, *α*_λ_ = 0.5, *μ*_0_ = 0.05, and *α*_*μ*_ = 0.1; (4) λ_0_ = 0.075, *α*_λ_ = 0.5, *μ*_0_ = 0.05, and *α*_*μ*_ = *-*0.4, -0.15, 0.35, 0.5. Values were chosen to test the most amount of variation in the λ and *μ* dependency, while reliably simulating surviving trees. For each combination of parameter values, we simulated 5000 trees using the RPANDA function sim_env_bd, stopping the simulations when the equivalent of 52 Myrs was achieved. The median tree size (of trees with more than 50 tips) was 280 for trees with a λ-dependency on T, 435 for trees with a *μ*-dependency on T, and 812 for trees with both a λ- and *μ*-dependency on T. Finally, we fitted the generating model (exponential dependence of λ or *μ* on *T*) by maximum-likelihood using the RPANDA function fit_env (53) and recovered parameter estimates for λ_0_, *α*_λ_, *μ*_0_, and *α*_*μ*_. The likelihood implemented in this function is the one described in (54) and (19); here we conditioned the likelihood on stem age. Throughout, we used a Nelder-Mead optimization algorithm (56) to fit models with a dependence of λ on time or temperature and a simulated-annealing optimization algorithm (6) to fit models with a dependence of *μ* on time or temperature. This is because for *α >* 0.1 in the latter models the Nelder-Mead algorithm did not converge – specifically, the optimization function could not be computed at the initial parameters for a wide range of initial parameters.

Because many empirical phylogenies are undersampled, we tested the effect of undersampling on inferring parameters in temperature-dependent trees. We simulated 1000 temperature-dependent λ trees (λ(*t*) = 0.2*e*^0^.^4*T*^ ^(*t*)^) and constant extinction (*μ* = 0.05) and 1000 temperature-dependent *μ* trees (*μ*(*t*) = 0.05*e*^0.1*T*(*t*)^) and constant speciation (λ = 0.5). We jackknifed the trees (i.e., sampled without replacement) at 10% intervals from 90 *-* 50% and inferred parameter estimates by fitting a temperature-dependent speciation model to each set of 1000 trees using the function fit_env,while accounting for undersampling by specifying the corresponding sampling fraction (54; (19). The median tree size (of trees with more than 50 tips) of the original temperature-dependent λ trees was 1209 tips and of the original temperature-dependent *μ* trees was 1119 tips.

We tested the ability to detect temperature-dependent diversification when it occurs and to not detect it when it does not occur. Many paleoenvironmental curves, despite fluctuations in their trends, sport a general tendency to increase or decrease over time.Therefore, in addition to testing the ability to distinguish between temperature-dependent and null, constant rate models, we tested the ability to distinguish between temperature-dependent and time-dependent models. We simulated birth-death trees with a positive and negative exponential dependency of λ on temperature (λ = 0.2*e±*^0.9*T*^, *μ*(*t*) = 0.05), with an increasing and decreasing dependency of λ on time (λ = 0.2*e±*^0^.^2*t*^, *μ*(*t*) = 0.05), and with constant rates (λ = 0.2, *μ* = 0.02). Additionally, because unaccounted-for background variation in diversification rates across lineages can potentially lead to biased inferences (64; 54; 66), we tested whether rates that vary among lineages can be misleadingly interpreted as environmental dependence. We therefore simulated trees with a 10% probability of a rate-shift in λ at each branching event, with an initial λ_0_ = 0.2; at each event where a shift occurred, the new λ was drawn from a normal distribution of values (with a standard deviation of 0.1) centered on the ancestral value; *μ* was held constant at 0.02. For each scenario we simulated 5000 trees; the median tree size (of trees with more than 50 tips) was 455 tips. Finally, to test whether high relative extinction rates affect our ability to properly infer temperature-dependence on speciation rates, we simulated 1000 temperature-dependent λ trees (median tree size of 190 tips) with λ = 0.75*e*^0.35*T*(*t*)^ and *μ* = 0.7. Time-dependent trees were simulated using tess.sim.age (38) and trees with rate-shifts were simulated with our own code. We fit all three models (exponential dependency on temperature, exponential dependency on time, and constant rates) to each tree and compared support using the corrected Akaike Information Criterion (AICc) (11). Fits of constant and time-dependent models were performed using the fit_bd function in RPANDA. We then calculated the percentage of each set of trees best fit by each model (the model with lowest AICc was selected). We likewise simulated birth-death trees with a fixed λ and a positive and negative exponential dependency of *μ* on temperature (λ_0_ = 0.4, *μ*_0_ = 0.1, *α*_*μ*_ = *±*0.4), an increasing and decreasing dependency of *μ* on time (λ_0_ = 0.5, *μ*_0_ = 0.01, *α*_*μ*_ = *±*0.4), and with no dependency of *μ* on time (i.e., constant λ_0_ = 0.2, *μ*_0_ = 0.01). For each scenario we simulated 5000 trees; the median tree size (of trees with more than 50 tips) was 358 tips. Here, in addition to fitting the corresponding models, we also fit models with an exponential dependency of λ on temperature. This was done to see whether a *μ*-dependency on temperature left a detectable signal in λ.

In order to have an idea of the effect of tree size on parameter estimation and the ability to recover temperature dependency, we simulated 5000 temperature-dependent λ trees (λ = 0.2^*e±*0.05*T*(*t*)^ and *μ* = 0.05). We fitted the corresponding temperature-dependent model and recovered parameter estimates for each tree as above. We also fitted the exponential time-dependent and constant rate models and computed the Akaike weights of these models on each tree. Finally, we reported the results by tree size bin. This procedure can potentially introduce a bias, because trees falling in a particular size-bin are trees that, by chance, even though they share the same simulated parameter values, diversified more (or less) than others. We did not observe such a bias, however. We similarly tested the effect of tree size on parameter estimation and model recovery for temperature-dependent *μ* trees (*μ* = 0.2*e*^0.1*T*(*t*)^); in these simulations, we varied λ_0_ across simulations so as to avoid the bias mentioned above. λ_0_ values were 0.1, 0.2, 0.3, 0.4, and 0.5.

There will necessarily be a loss of resolution between the actual fluctuations in an environmental variable over time and the reconstruction of those fluctuations with sampled data. It is therefore important to understand how sensitive the environment-dependent model is to the reconstructed fluctuations. To evaluate this, we tested the effect of discrepancies in resolution between the temperature curve used to simulate trees and the one used to fit the temperature-dependent model. We simulated pure-birth (*μ* = 0) trees with an exponential dependency between λ and temperature, where the smoothing function for the temperature curve was determined using generalized cross-validation (34). We simulated 5000 trees with positive temperature-dependency (λ = 0.2*e*^0.05*T*(*t*)^) and 5000 trees with negative temperature-dependency (λ = 0.2*e*^-0.05*T*(*t*)^). The median tree size (of trees with more than 50 tips) was 610 tips. We then fitted temperature-dependent models to the simulated trees, using increasingly smoothed temperature curves. Smoothed curves are meant to represent degraded environmental data and were obtained by reducing the number of degrees of freedom (DOFs) applied to the smoothing function. For each tree, we compared parameter estimates, as well as statistical support in comparison with time-dependent models (with exponentially trending λ) obtained when fitting temperature-dependent models with increasingly smoothed temperature curves.

### Performance of model selection on trees simulated with various environmental dependencies

Generalizing the analyses performed for temperature-dependency, we assessed whether environmental dependence can be detected on trees simulated with λ dependencies on specific paleoenvironment curves. Because we did not see any effect of extinction on the ability to recover the environment-dependent model in the case of temperature-dependency, we simulated pure-birth trees (*μ* = 0) with exponential dependencies between λ and each paleoenvironmental curve X (see *Paleodata* above). We simulated trees at various values of *α*_λ_ (0.15, 0.3,0.45, 0.6, 0.75, 0.9, 1.05, 1.3, 1.6, 1.9, 2.2) to test how the strength of the simulated dependency influenced the recoverability of the environmental model. For each value of *α*_λ_ and for each paleoenvironmental curve we simulated 1000 trees with a median tree size (for trees with more than 50 tips) of 310 tips. λ_0_ was identical across paleoenvironmental curves, but varied between *α*_λ_ values, in order to maximize the number of surviving, reasonably sized simulated trees. For each tree, we fitted the same environment-dependent model that was used to simulate that particular tree, as well as a constant-rate λ model and a time-dependent λ model, and compared their statistical support with AICc. We selected the best model as the one with the lowest AICc support. We also compared the relative support of the models using Akaike weights.

We carried out the same type of analyses to assess whether an environmental dependence on *μ* can properly be inferred. We simulated trees with an exponential dependency between *μ* and a paleoenvironmental curve X, holding λ constant, for different values of *α*_*μ*_ as above. For each value of *α*_*μ*_ and for each paleoenvironmental curve we simulated 1000 trees; the median tree size (of trees with more than 50 tips) was 320. For each tree, we fitted the same environment-dependent model that was used to simulate that particular tree 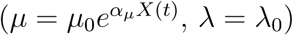 as well as a constant-rate birth-death model and a model with *μ* exponentially dependent on time and constant λ. We also fitted a model where the environment influences speciation rather than extinction rates 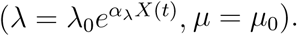. We did this under the hypothesis that trees simulated under a *μ* dependency on a paleoenvironmental curve could manifest a signature of that dependency in their λ.

Environmental curves differ widely in their shape and characteristics (Fig 3B). In particular, the biotic curves that we considered here (constructed from fossil data) appeared coarser than the abiotic ones. To assess whether environmental dependency was more easily detected for certain types of curves than others, we characterized the curves based on defining attributes, such as biotic/abiotic, linearity/non linearity, and six further metrics, two capturing temporal autocorrelation, two capturing global trend, and two capturing the composition of the data. Next, we assessed the effect, if any, these features had on the rate of recovery of the models at different values of *α*_λ_. To test for linearity, we tested for the presence of one or two breakpoints in the data using the threshold test (36). We used two metrics reflecting temporal autocorrelation; first we measured the correlation between values in the original time-series and a new series that lags behind by some amount of time, for a series of time lags. We then fit regressions between the various time lags and the corresponding correlation value. We used the intercept of the regression to measure short-term autocorrelation, and the slope to measure the rate at which autocorrelation decreases. A global trend analysis was conducted by fitting a linear model using generalized least squares to each curve and computing the correlation, slope, and degrees of freedom for the fitted model. Finally, we estimated the average rate of change of the curves by calculating the mean over all time periods of the slope between values within each time period.

Finally, for trees simulated under each paleoenvironmental curve with an exponential dependency on λ, we compared fits for models using all the paleoenvironmental curves. For each set of simulated trees, we calculated the percentage of trees best fit (i.e., lowest AICc) by each paleoenvironmental model and reported this percentage as a function of *α*.

### Testing macroevolutionary hypotheses of environment-dependency on empirical phylogenies

We used time-calibrated molecular trees for Cetacea, Ruminantia, and Portulacaceae. The Cetacean phylogeny was 98% sampled and constructed from six mitochondrial and nine nuclear genes in a Bayesian framework; it was time-calibrated using fossil data; and divergence dates were estimated using a relaxed molecular clock (78). For Ruminantia we used two different trees: one was constructed using full mitochondrial genomes and 16 fossil calibration points in a Bayesian framework, but was only 65% sampled (9); the other one was constructed using a supermatrix of 124 previously published trees and was 100% sampled (12). The two phylogenies differed markedly in their datation; in particular, the phylogeny of (9) hypothesized significantly more recent crown ages for many families and for Ruminantia than that of (12). For Portulacaceae, the tree was 40% sampled and constructed separately using a combined matrix of genomic markers for chloroplast in a maximum likelihood framework and Bayesian inference; divergence times were estimated using a relaxed clock calibrated on relevant geological events.(58). We conducted analyses on Bayesian posterior trees for Cetacea and both Ruminantia trees; we analysed both the maximum-likelihood and Bayesian tree for Portulacaceae. The above trees were chosen because diversification in each has previously been associated with an abiotic process(78; 58; 12).

The nine paleoenvironment models plus a constant-rate birth-death model and time-dependent model (with and without extinction) were fit to the empirical phylogenies, accounting for missing taxa by applying the relevant sampling fraction (54; (19). For each paleoenvironment model, we tested λ and *μ* as constant, linear, and exponential functions of the paleoenvironmental curves, as well as *μ* = 0. The best-fit model for each phylogeny was determined by Δ*AICc* as above, and the corresponding speciation and extinction parameter estimates were recorded. All palaeoenvironmental curves were tested. While some variables (temperature and sea level) are directly biologically relevant for the three clades, other variables (*CO*_2_, *δ*_13_*C*, silica, archaeplastida diversity, ostracod diversity) are directly relevant for some clades but not all, and others still (foraminifera and radiolarian diversity) are not directly biologically relevant to any of the clades. When environmental variables were not directly relevant, we still included them as negative controls. We also note that while significant support for environment-dependent models is typically interpreted by an effect of the environment on clade dynamics, such support could also occur if the rise or demise of a given clade impacts the targeted environmental variable (e.g. silica levels in the ocean are often attributed to the expansion of land plants). Hence, environmental variables that are biologically relevant for a given clade are those that can either influence, or be influenced by, that clade.

All clades can potentially be influenced by temperature and sea level changes. For cetaceans, aside from these two variables, we may expect a dependency on silica levels, archaeplastida, and ostracods. Indeed, the evolution of cetaceans has been influenced by diatoms, which are phytoplankton that require silica (48). Archaeplastida include green algae that can have a negative impact on diatom blooming (41), and thus potentially also affect cetaceans. Finally, ostracods may also be expected to influence cetacean diversification, either directly as a food source or indirectly as diatom predators. For Portulacaceae, biologically relevant variables are *CO*_2_, *δ*_13_*C*, silica, and Archaeplastida. Indeed, reductions in *CO*_2_ levels have been linked to radiations of angiosperms during the Cretaceous and the rise of *C*_4_ plants, such as Portulacaceae, during the Miocene (33), which is also reflected in global levels of *δ*^13^*C*. Finally, Archaeplastida include land plants that can engage in mutualistic or competitive interactions with Portulacaceae. For ruminants, changes in plant life across the globe have been crucial, such that all variables mentioned for Portulacaceae (*CO*_2_, *δ*_13_*C*, silica, and Archaeplastida) are also relevant through their presumed effect on land plants. Archaeplastida can in addition directly influence diversification in Ruminantia via change in their feeding regime.

In order to illustrate how one could test the relative importance of abiotic *versus* biotic variables, we selected the best supported abiotic and biotic variables and computed the relative probability of these two models based on their Akaike weights; we also computed the Akaike weight of each model among the set of all models and summed these Akaike weights over all the abiotic variables (or biotic variables) to obtain an estimate of the overall support for abiotic (or biotic) variables. The first approach has the advantage to not be biased by the total number of variables, or biologically relevant variables, in each category (abiotic and biotic). However, it measures the relative importance of the two most supported abiotic and biotic variables rather than the relative importance of abiotic and biotic variables as a whole. Intermediate approaches could be envisioned where the relative support for abiotic and biotic variables is computed based on a fixed, homogenized number of biologically relevant biotic and abiotic variables.

Additionally, for each phylogeny, we assessed the support of the best-fit paleoenvironmental model with smoothing splines computed with increasingly smaller DOFs. This was done in order to test at what resolution of the paleoenvironmental curve support for environment-dependent diversification started to be lost. We compared all fits using AICc as above.

## Results

### Performance of temperature-dependent diversification models

Temperature-dependent models were able to accurately recover simulated parameter values. Maximum-likelihood estimates of the parameters of the temperature-dependent λ models were unbiased (Figure 1), even when extinction rates were high (Figure 1D). Maximum-likelihood estimates of the parameters of the temperature-dependent *μ* models were likewise unbiased, but they were more variable (Supplemental Figure 1). When both λ and *μ* were exponentially dependent on temperature, parameter recovery remained reliable (Supplemental Figure 2).

**Figure 1:**
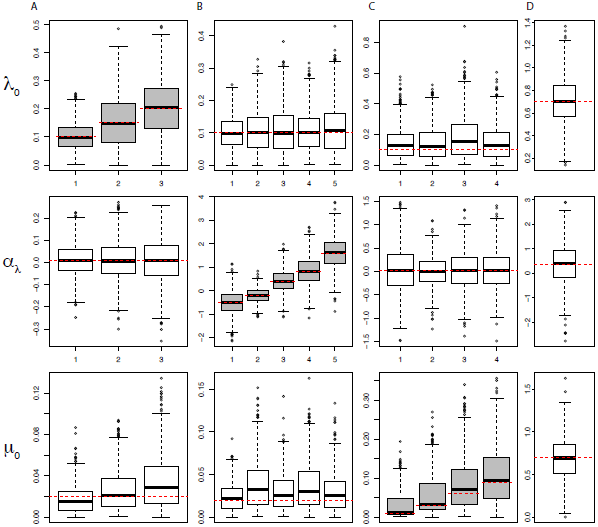
(A) Recovered parameter estimates for trees simulated with an exponential dependency of λ on temperature. Simulations with (A) varying λ_0_ and constant *α*_λ_ and *μ*_0_, (B) varying *α*_λ_ and constant λ_0_ and *μ*_0_, and (C,D) varying *μ*_0_ and constant λ_0_ and *α*_λ_. Dashed red lines mark the simulated parameter value.

**Figure 2:**
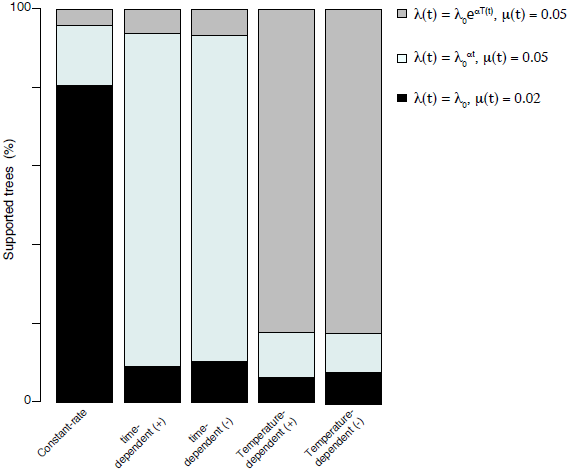
The ability to correctly recover trees simulated under a constant-rate model,positive and negative dependence of λ on time models, and positive and negative dependenceof λ on temperature models. Each column shows the percentage of trees simulated under themodel specified on the x-axis that finds support for each model as specified in the legend.

Undersampling did not bias parameter estimates, but affected the uncertainty around the estimates (Supplemental Figure 3). While the median estimates did not deviate significantly from the simulated estimates (*P <* 0.01), the variance increased with decreasing sampling fraction.

**Figure 3:**
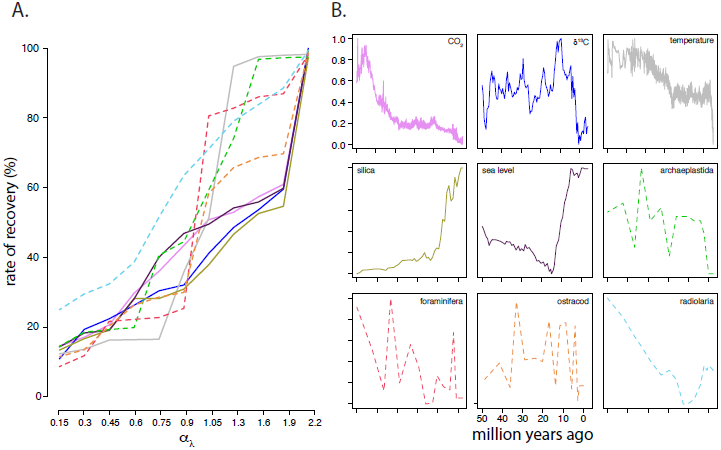
Model identifiability for trees simulated with an exponential dependency of λ on different paleoenvironments, X, (λ= λ_0_*e*^α*X(t)*^ and μ = 0) for varying values of *α λ* (A) Rateof recovery is defined as the percentage of trees with better AICc support for the simulating environment-dependency model versus an exponential time-dependent model and constant rate model. (B) Scaled paleoenvironmental curves. Colors in (A) and (B) correspond. Dashed lines correspond to biotic variables.

Temperature-dependent speciation models can be correctly distinguished from other models. Each model – constant, time-dependent, and temperature-dependent λ models – were overwhelmingly recovered when they were the generating model (*≥* 75%, Figure 2), even when relative extinction was high (Supplemental Figure 4). Trees simulated with a time-dependency on λ were rarely (*~* 5%) recovered by a temperature-dependent model and *vice versa*, which is especially significant given the overall tendency for the temperature curve to decrease with time. Heterogeneity in speciation rates across lineages did not lead to a false support for temperature dependency: trees with speciation rate shifts were recovered by constant-rate models 32% of the time, models with increasing speciation rate through time 62% of the time, and temperature-dependent models only 6% of the time. The support for increasing speciation rate models can be explained by the fact that lineages that by chance have high speciation rates leave more descendants, such that the average speciation rate over lineages increases through time. The ability to detect temperature-dependency on *μ* when it occurred was considerably lower: *~* 20% of trees simulated with a temperature-dependency on *μ* were accurately recovered by the generating model (Supplemental Figure 5).

**Figure 4:**
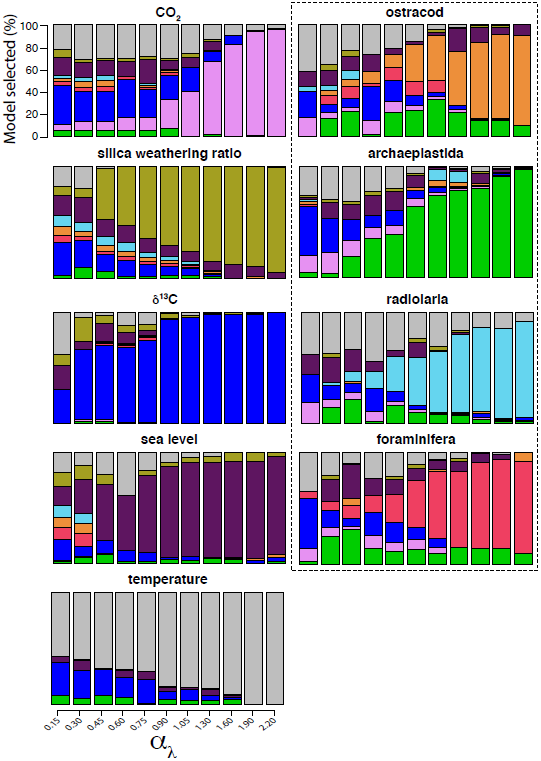
Rate of model identifiability between environment-dependent models. For each set of trees simulated with an exponential dependency of λ on a particular paleoenvironmentalcurve, X (λ= λ_0_*e*^αX(t)^ *μ*= 0), the percentage of trees recovered by each paleoenvironmental model for diffierent values of *α λ*. The simulated model is listed above each barplot. Colors correspond to Figure 3; abiotic variables are circumscribed by the dashed box.

**Figure 5:**
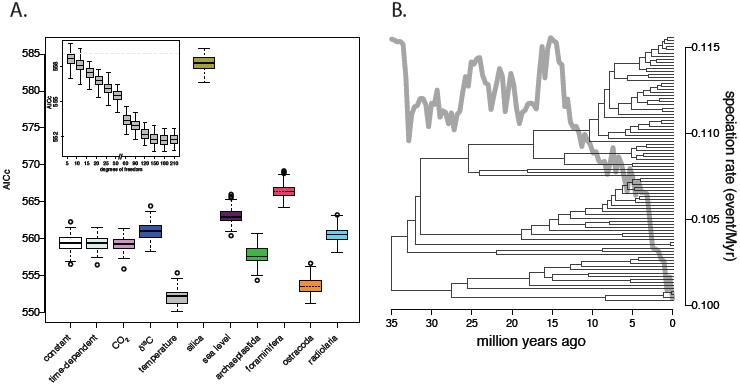
Environment-dependency in Cetaceans. (A) AICc support for different environment-dependent models, a constant-rate birth-death model, and an exponential time-dependent model (without extinction) on a distribution of 100 posteriorly sampled probabilities of the Cetacean phylogeny. Colors and line type correspond to Figure 3. (A, inset) AICc support for temperature-dependent models fitted with smoothed splines defined by varying DOFs. AICc value for the time-dependent model is shown (dashed line). (B) The consensus Cetacean phylogeny and over time inferred from the temperature-dependent model (on the consensus tree).

As expected, the ability to recover parameter estimates and the temperature-dependent model improved with tree size. For temperature-dependent λ models, median parameter estimates approximated simulated values precisely as soon as trees had more than 100 tips, and the variance of recovered estimates decreased considerably with tree size (Supplemental Figure 6). With the *α*_λ_ values considered in these simulations (*α* = *±*0.05), the Akaike weight was well above 0.5 for trees with more than 200 tips, and steadily increased to 0.9 for trees with more than 900 tips (Supplemental Figure 6). For temperature-dependent *μ* models, median estimates for *α*_*μ*_ approximated simulated values for trees with more than 400 tips and the range of estimates decreased considerably for trees with more than 600 tips (Supplemental Figure 7). Median estimates for *μ*_0_ were slightly higher than simulated values for all tree sizes, which is consistent with our previous estimates of *μ*_0_ at *α ≥* 0.1 (see Supplemental Figure 1). The Akaike weight of the temperature-dependent *μ* model increased with tree size, but in agreement with the results above it remained low, below 0.4 even for trees with more than 800 tips (Supplemental Figure 7).

**Figure 6:**
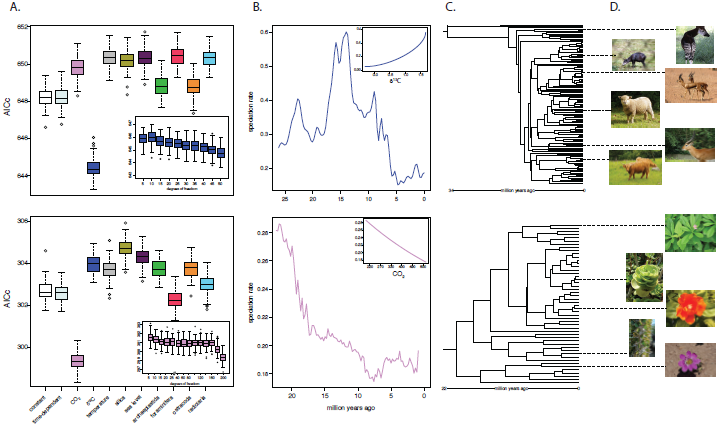
Diversification dynamics in (top) Ruminantia and (bottom) Portulacaceae. (A) AICc support for different environment-dependent models, a constant-rate birth-death model, and an exponential time-dependent model (without extinction). (A, inset) AICc support for the best supported model fitted with smoothed splines defined by varying DOFs. (B) λ over time inferred by the best supported model in (A). (B, inset) λ as a function of the environment for the best fit model, where λ is shown as a function of a non-scaled environmental curve. (C) Consensus phylogeny and (D) representative organisms: (top to bottom) *Okapia johnstoni, Cephalophus silvicultor, Gazella gazella, Ovis aries, Kobus leche, Bos taurus, Talinum paniculatum, Portulacam molokiniensis, Pereskia sacharosa, Alluaudia ascendens, Montiopsis andicola. The figures are available here (top to bottom):* https://www.ickr.com/photos/dkeats/5269538538/sizes/z/, https://www.ickr.com/photos/7326810@N08/1332824505,https://www.ickr.com/photos/93882360@N07/12801059065,https://commons.wikimedia.org/wiki/User:AnemoneProjectors,http://www.naturfakta.no/dyr/?id=1809,https://www.ickr.com/photos/computerhotline/60011372369_Portulaca_molokiniensis.jpg,https://www.ickr.com/photos/jimmytst/11078000494/in/photolist-hSVDAw-abxoPu,https://www.ickr.com/photos/junglemama/5364129725,http://www.nhm.ac.uk/natureplus/blogs/solanacae-in-south-america/tags/argentina.

When fitting models with degraded temperature curves, we found that accurate parameter estimates could still be properly recovered, unless the environmental data were very highly degraded (i.e., for a smoothed spline constructed with *≤* 15% of the total DOFs available in generalized cross-validated model (Supplemental Figure 8). Fitting models with a degraded temperature curve slightly reduced the ability to distinguish temperature-dependent models from time-dependent ones, but this ability remained good, with *<* 15% of the phylogenies simulated under a temperature-dependent model with the most degraded curve (DOFs= 5% of the generalized cross-validated value) finding support for time-dependent rather than temperature-dependent models (Supplemental Figure 8).

### Performance of model selection on trees simulated with various environmental dependencies

The ability to detect environment-dependence when it exists does not drastically change with the paleoenvironmental variable considered. For all nine paleoenvironmental variables, environment-dependent speciation can be detected and distinguished from a constant rate and time-dependent effect, as soon as the environmental effect is relatively strong (Figure 3). The ability to distinguish between the simulating environment-dependent model and a pure-birth or time-dependent model increases roughly linearly with *α*_λ_, from approximately 20% at *α*_λ_ = 0.15 to 100% at *α*_λ_ = 2.2. This result is unlikely to be an effect of tree size, as the median tree size does not vary much across *α*_λ_ values (185 *±* 36). As *α*_λ_ increases, so does the support for the simulated environmental model, as defined by the Akaike weight support for the environment-dependent, time-dependent, and constant-rate models (Supplemental Figure 9), consistent with expectations. For *α*_λ_ *>* 1, curves for biotic palaeodata, which tend to be coarser than abiotic curves, show, on average, higher recovery rates, while for *α*_λ_ *<* 1, curves for abiotic palaeodata show, on average, higher recovery rates (Supplemental Figure 10A). There is no such noticeable effect on Akaike weights, though (Supplemental Figure 9). There is no apparent effect of linearity, the level of autocorrelation, the direction or magnitude of the best-fit regression slope, degrees of freedom, or the average rate of change of the environmental curves on our ability to recover how the corresponding environment affected speciation (Supplemental Figure 10B-C).

Environment-dependent extinction is generally not detected, regardless of the strength of the environmental effect: the ability to recover environment-dependent *μ* is *<* 10% across all *α*_*μ*_ values (Supplemental Figure 11). Even when only trees with more than 400 tips are considered (median tree size = 1183 tips), the ability to recover the correct model does not exceed 20%. Environment-dependent extinction is more often detected as an environmental dependence on λ than on *μ* (Supplemental Figure 11), but this still happens in less than 20% of the trees.

The ability to distinguish between λ-dependence on different environmental variables is good as soon as the environmental dependence is strong enough: while a considerable percentage of trees were recovered by the wrong environment-dependent model for low *α*_λ_ (a condition under which the environmental model would likely not be selected when compared to a time-dependent model (Figure 3), the correct model was recovered in the majority of trees for *α*_λ_ *>* 1 (Figure 4). For several paleoenvironmental dependencies – specifically, *δ*^13^*C*, sea level, and temperature – the simulating model was recovered the majority of the time, even at low *α*_λ_ values. These three environmental variables were also those that were most often misleadingly detected for trees generated under other environmental dependencies (Figure 4). As above, the median tree size did not vary much across *α*_λ_ values (249 *±* 34).

### Testing macroevolutionary hypotheses of environment-dependency on empirical phylogenies

The Cetacean phylogeny was best fit by a pure-birth exponential speciation model with a positive dependency on temperature across all posterior distributions (λ = 0.097*e*^0.02*T*(*t*)^ for the consensus phylogeny) (Figure 5). The probability that the abiotic model is the best, as calculated by the Akaike weight for the best-fit abiotic model (temperature) when compared to the best-fit biotic model (ostracod diversity) was 0.72. The sum of Akaike weights across abiotic variables was 0.73 *±* 0.12 and across biotic variables 0.27 *±* 0.15. We furthermore compared fits of the temperature-dependent λ model with splines smoothed by a range of DOFs (5 *-* 190). Using the maximum DOFs (208), as defined by generalized cross-validation, the temperature-dependent model was significantly better supported than a time-dependent model (Δ*AICc* = 6.9) and other environmental models (Δ*AICc >* 1.3). The time-dependent model only showed a comparable fit to the tree when the temperature curve was smoothed with DOFs below 10. This suggests that diversification in Cetaceans is best supported by the global trend in the temperature curve, although the decrease in AICc support with reduced DOFs is evidence that diversification is also supported by specific fluctuations within the temperature curve. The Ruminantia phylogeny of (9) was best fit by a pure-birth exponential speciation model with a positive dependency on *δ*^13^*C* across all posterior distributions (λ = 0.175*e*^0^.^7^*δ*13*C* for the consensus phylogeny) with an Akaike weight for that model (*δ*^13^*C*) when compared to the best-fit biotic model (ostracod diversity) of 0.94 (Figure 6A). The sum of akaike weights across abiotic variables was 0.79 *±* 0.06 and across biotic variables 0.11 *±* 0.06. The Ruminantia supertree of (12), however, was best supported by a constant-rate birth-death model (λ = 0.11, *μ* = 1.20*e -* 7) (Supplemental Figure 12). The analysis by (12) separated dietary groups and found maximum support for a model with a linear dependency of temperature on λ; in our analysis (which did not separate dietary groups), a model with linear dependency on temperature found considerably poorer support than the constant-rate model (Δ*AICc* = 142.329). Portulacaceae was best fit by a pure-birth exponential λ model with a negative dependency on *CO*_2_ for both the maximum-likelihood and Bayesian phylogeny (λ = 2.98*e*^-0^.^44^*CO*2 for the maximum-likelihood phylogeny) (Figure 6B), with an Akaike weight for that model (*CO*_2_) when compared to the best-fit biotic model (foraminifera diversity) of 0.82. The sum of Akaike weights across abiotic variables was *±* 0.04 and across biotic variables 0.25 *±* 0.04. When the curves of the best-fit models for Ruminantia (using the phylogeny of (9)) and Portulacaceae were smoothed with increasingly fewer DOFs and refit to the data, there was little effect on the computed AICc for Ruminantia, but a steep increase in AICc values for Portulacaceae at DOFs below 90%, and then again at DOFs below 80% of total DOFs, suggesting that diversification in Ruminantia is supported by the general trend of the *δ*^13^*C* curve, whereas in Portulacaceae it is supported by both the general trend and minor fluctuations of the *CO*_2_ curve. For both phylogenies, AICc support for the best-fit environment-dependent model became poorer than for a time-dependent model only when the DOFs dropped to 2, which effectively made the environment-dependent models equivalent to time-dependent models.

## Discussion

The history of Earth has been marked by shifting land masses, volcanic eruptions, and continuous global atmospheric change, as well as a widespread and often dramatic turnover and dispersal of life. There is general recognition that changes to biodiversity over geological time are driven by the environment (26) and species interactions (81). The proportional contribution of various abiotic and biotic drivers within individual clades and across the tree of life, however, is still debated (30). This is due, in part, to the challenges of estimating the effects of these abiotic and biotic factors on clade diversification. We have tested a phylogenetic approach to address these challenges under a model-based framework that allows us to directly relate speciation and extinction rates in a clade to biotic and abiotic factors in the paleoenvironment. Applying this modeling framework to empirical phylogenies, we identify specific environmental variables that likely played a major role in shaping the evolution of the corresponding clades.

Our simulation results point to three aspects of the use of environment-dependent models that should be considered. Firstly, we show that speciation and extinction parameter estimates do not systematically deviate from true values as undersampling increases (as long as undersampling is accounted for when fitting the models), although the variability of estimates certainly does. Secondly, we find that the accuracy of model selection depends strongly on the strength of the environmental dependence measured by the *α*_λ_ parameter (effect size) and tree size. Finally, we find notable differences in the ability to recover environmental dependency across distinct paleoenvironmental curves. In particular, in the set of paleoenvironmental curves that we considered, recovering the proper environmental dependence under strong dependence was easier for the biotic variables and temperature in comparison with the other abiotic variables. We could not, however, characterize which features of the time-series data influence its recovery rate. A thorough understanding of how differences in the shape of the paleoenvironmental curves influence our ability to make inferences will require simulating environmental curves with distinct and controlled characteristics.

It has been noted before that, although possible in theory, inferring extinction rates from molecular phylogenies is difficult in practice (see (64; 52; 4; 65)). Inferring the environmental dependence of extinction rates is, not surprisingly, also difficult. Specifically, we find that inferring extinction parameters is very variable for small trees (*<* 200 tips) and that accurately distinguishing *μ*-dependent environmental models from constant-rate or time-dependent models can be unreliable.

The strengths and limits of the environmental models identified here suggest some guidelines for best practice. First, parameter estimates for environment-dependent speciation models start to be accurate for trees with more than 100 tips (although this number, of course, also depends on effect size and the nature of the curve). Second, all results related to environmental dependence in extinction should be treated with care, especially for trees with fewer than 400 tips. Second, environment-dependencies that may not seem biologically relevant can be useful negative controls or used to detect potential hidden variables. In our analyses, we did not find strong support for any biologically irrelevant variable, which gives us confidence in the dependencies for which we did find strong support. Third, if a user is interested in analyzing the effect of an environmental variable not considered here, we recommend that s/he directly analyses the statistical properties of the environment-dependent model applied to that particular variable using simulations. Finally, when testing environmental dependencies in speciation rates, inferred *α*_λ_ values for the best and second-best fitting models can be consulted; if there is a high false discovery rate for the best-fit model at the *α*_λ_ value inferred for the second-best-fit model (see Figure 4), then the result should be interpreted cautiously.

A recent work that used the temperature-dependent model (19) to infer Diversification in modern birds found rates of speciation and extinction to be negatively dependent on temperature (16). We can assess how robust this result is in light of our simulation analyses. The temperature curve was smoothed with *~* 25% of the DOFs inferred by generalized cross-validation. Given our observation that the temperature-dependent model is robust to smoothing, we expect this smoothed curve to be sufficient to generate accurate parameter estimates. The tree has 230 tips, which is sufficient for accurately inferring parameter values for λ, but not necessarily for *μ*. The result that extinction is temperature-dependent, therefore, should be interpreted with caution. The tree is severely undersampled, with a sampling fraction of *~* 0.02%, which might produce estimates for both speciation and extinction considerably deviated from their true values. Finally, the temperature-dependent model-fitting was conducted on only one tree (the maximum clade credibility tree), which is not ideal. Dating discrepancies can be sources of potential discordance between results, which is something we account for in the Cetacean and (both) Ruminantia phylogenies by analyzing posteriorly sampled trees.

In agreement with previous macroevolutionary (19; (48) and macroecological (86) work, we find that diversification in Cetaceans was positively associated with temperature. The best-fit model is a pure-birth model with exponential dependence of λ on temperature. We show in addition that temperature-dependence is better supported than dependence on other paleoenvironmental variables. It has been suggested that Plio-Pleistocene fluctuations in oceanic temperatures created ecological opportunities for allopatric speciation in delphinids (78) and this may be why we find diversification in Cetaceans to be dependent on temperature. Notably, the inferred *α*_λ_ value for the second-best-fit model, ostracod diversity, is 0.27 and, at this *α*_λ_ value, temperature shows a relatively high false discovery rate (around 40%). Therefore, the temperature-dependent result should be interpreted cautiously. The support we find for a positive dependency of ostracod diversity on diversification in Cetaceans (λ = 0.089 *\* e*^0^.^193^*\*ostracoddiversity*) suggests that the role of ostracods as a food source may have positively affected Cetacean speciation.

In Ruminantia, the results depended on the phylogeny we used, which is not surprising given the markedly different datations hypothesized by these two phylogenies. With the phylogeny of (9), we found diversification to be best supported by a pure-birth exponential speciation model with a positive dependency on *δ*^13^*C*. This substantiates a major role for *C*_4_ grasses, which became dominant during the Miocene and have been hypothesized to have spurred diversification in ruminant mammals (12). The *δ*^13^*C* data we used were collected from marine samples rather than from terrestrial samples (e.g., bovid tooth enamel), and so are not a direct measurement of the relative abundance of *C*_4_ and *C*_3_ grasses. However, there are notable similarities in these data, particularly during the Neogene (13). For the Ruminantia supertree of (12), we find support for a constant-rate birth-deal model rather than any environment-dependent model, which is contrary to what was found in (12). This may be because that study conducted separate analyses on different dietary groups.

Our analysis of Portulacaceae, which was best fit by a pure-birth model with a negative exponential dependency on *CO*_2_, adds statistical evidence to the often described macroevolutionary relationship between *CO*_2_ levels and angiosperm diversification.

Previous work has shown that angiosperm diversity in the Cretaceous was negatively correlated with atmospheric *CO*_2_ (23). The reason for this negative correlation, however, is debated: some argue that so-called “*CO*_2_ starvation” drove radiations in angiosperms as they competed and searched for new niches (79; (69); others that diversification in angiosperms caused a decline in *CO*_2_ through the geological weathering of calcium and silica (43; (82). Our result cannot resolve this conflict, but does extend the previously observed negative relationship into the Cenozoic for an angiosperm family, and furthermore quantifies the effect of *CO*_2_ levels on that family‘s diversification. We can be confident that the *CO*_2_ model is not a false positive, as *CO*_2_ is rarely selected when it is not the generating model (Figure 4). In particular, when the second-best-fit model (foraminifera diversity) is the generating model, there is a zero or near-zero selection for the *CO*_2_ model, even at much lower *α*_λ_ values than the one estimated here (4.47) for the dependency to foraminifera (Figure 4).

For all three phylogenies, we estimated the support of all biotic and abiotic models, regardless of whether they were biologically relevant or not. In the three clades, we found a strong support for the best abiotic model when compared to the best biotic one, as well as a stronger support for the all abiotic variables combined compared to all biotic variables combined. This result may be biologically true, in which case it emphasizes the overarching importance of the global abiotic environment on species diversification in vastly different organisms. It may instead reflect the lack of relevant biotic variables we tested.

We note that the absence of extinction inferred in Cetacea and Ruminantia is not realistic, and is inconsistent with what we know from the fossil record (63; (12). Accounting for heterogeneity in diversification rates across these groups could help in recovering more meaningful extinction rate estimates (54). This could be done by applying the same type of tree partitioning that was used by (54), while fitting temperature-dependent models. Even though extinction seems to be poorly estimated in Cetacea and Ruminantia, this does not affect the confidence we can have in our inference of the dependency of speciation rates to environmental changes: our simulations suggest that high levels of extinction do not affect the ability to properly infer environment-dependent speciation, and that environment-dependent extinction is not mistakenly interpreted as environment-dependent speciation.

There are many potential extensions of the environment-dependent models presented here. First, paleoenvironmental variables not included in this work could be considered, such as planetary albedo or the origination rate of marine guilds. Second, models accounting for the simultaneous effect of several paleoenvironmental variables as Well as their interaction may provide more resolution on how various abiotic and/or biotic factors combine to influence species diversification. While fitting such models is already possible with the current implementation (any kind of dependence of speciation and extinction rates with environmental variables can be imputed, including dependencies on several environmental variables and their interactions), one would need to test the performance of such models using simulations before trusting the associated empirical results. Implementing the potential influence of hidden variables as well as a variable selection procedure would also be important directions for future work.

The way we have accounted for effects of the biotic environment is by considering interclade biotic interactions (e.g., the effect of ostracod diversity through time). However, there is a wide range of intraclade processes that are thought to influence diversification dynamics. For example, speciation may slow down as ecological opportunity recedes and interspecific competition intensifies following an adaptive radiation, resulting in a diversity-dependent effect (62; (27). Alternatively, speciation may accelerate if intra-clade species interactions (such as competition) promote reproductive isolation through phenotypic divergence (71; (2). Future forms of the environment-dependent model, therefore, could be developed to additively or synergistically account for intraclade and interclade effects. Similar models were recently developed for analyzing fossils (46; (29). Models directly incorporating intraclade effects on phenotypic divergence have also started to be developed (57; (22). Accommodating such factors in environment-dependent models would be useful for evaluating the relative effect of the environment, intra-and interclade species interactions, as well as their combined effect, on clade diversification.

We can also imagine adapting the model to analyse population genetic models on genealogical trees. Here, the environment-dependent model could be used to analyse how demographics (birth and death rates) have varied with time-dependent environmental variables (e.g., the rate of influx of immigrants in a region, the density of air pollution in a city). In traditional population genetic models, population structure is measured as a function of gene flow (88). Changes in gene flow can be the result of recent increases in population size or rates of dispersal; and so the factors that promote or limit gene flow cannot be easily inferred (74). The environment-dependent model, adapted to analyse the effects of, for example, habitat range contraction, environmental disturbance, or precipitous geographic isolation on birth and death rates within a population may therefore help resolve the contribution of those factors to changes in the population. As above, one would need to test the performance of such models using simulations before trusting the associated empirical results.

The phylogenetic comparative toolbox is being extended to better account for the effect of environmental changes. Similar to its use here, where we have shown the effect of different environmental variables on diversification in clades, we have seen a growing body of work relating, for example, decreases in temperature to accelerations in diversification in birds (16), increases in sea level to increased extinction in butterflies (21), and even orogeny to Andean plant diversification (61). Models of phenotypic evolution accounting for environmental variations have similarly been developed recently, and have found decreases in temperature to drive increases in the rate of body mass evolution in birds and mammals (17). Future analyses with these two types of models should allow us to better understand whether diversification and phenotypic evolution respond similarly (or differently) to past climatic changes, as well as the link between environmental variations, phenotypic divergence, and diversification.

Environment-dependent diversification models provide a statistical forum for testing a range of macroevolutionary hypotheses on species trees, implemented in user-friendly software. They provide a means for identifying which paleoenvironmental variable has most impacted diversification in a clade and how specifically that variable has impacted diversification. We have analyzed the statistical behavior of and provided guidelines for an environment-dependent model of species diversification. We think this will help researchers test fundamental macroevolutionary hypotheses on the effects of various abiotic and biotic factors on species diversification.

## Acknowledgments

We would like to thank Julien Clavel, Jonathan Drury, Guilhem Sommeria Klein, Marc Manceau, Cesar Martinez, and Olivier Missa for helpful comments on the manuscript, Odile Maliet for comments and her code to simulate rate-shifted phylogenies, and Juan Cantalapiedra for supplying the distribution of resolved Ruminantia supertrees. EL would like to thank Evan Charles for helpful discussion. Funding was provided through a European Research Council grant (ERC-CoG-PANDA) attributed to HM.

---

**Supplemental Figure 1:**
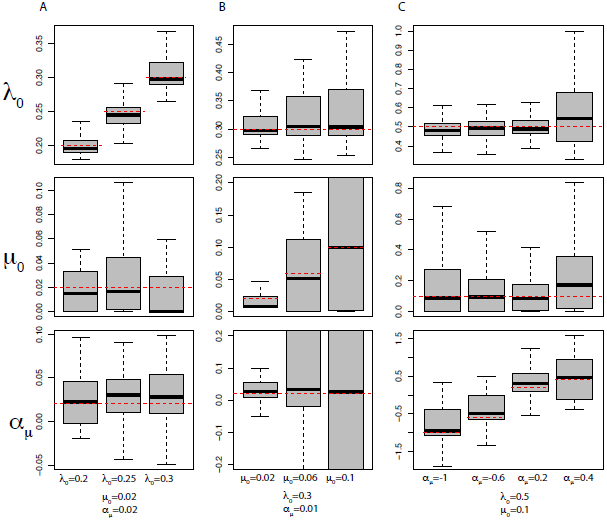
Recovered parameter estimates for trees simulated with a con-stant λ and an exponential dependency of μ on temperature. Simulations with: (A) varying λ_0_, constant μ_0_, and constant α_μ_; (B) constant λ_0_, varying μ_0_, and constant α_μ_; and (C) constant α_0_, constant μ_0_, and varying _μ_. Dashed red lines mark the simulated parameter value.

**Supplemental Figure 2:**
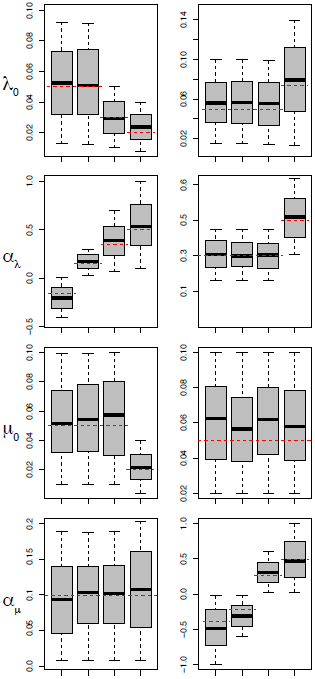
Recovered parameter estimates for trees simulated with an exponential dependency of λ and μ on temperature. Dashed red lines mark the simulated parameter value.

**Supplemental Figure 3:**
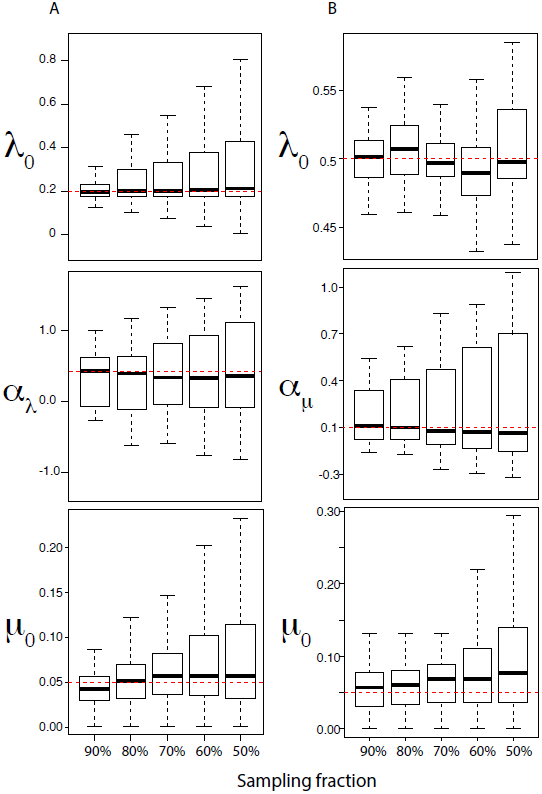
The effect of undersampling on parameter estimation. Parameter estimates for trees simulated with (A) a positive exponential dependency of λ on temperature and constant extinction (μ = 0.05) and (B) a positive exponential dependency of μon temperature with constant speciation (λ = 0.5) by fitting the temperature-dependent model. Parameter estimates are shown for trees with increasingly smaller sampling frac-tions, which were achieved by jackknifing the simulated trees by a fixed % and then fitting the environment-dependent model. Simulated parameters are marked by dashed red lines.

**Supplemental Figure 4:**
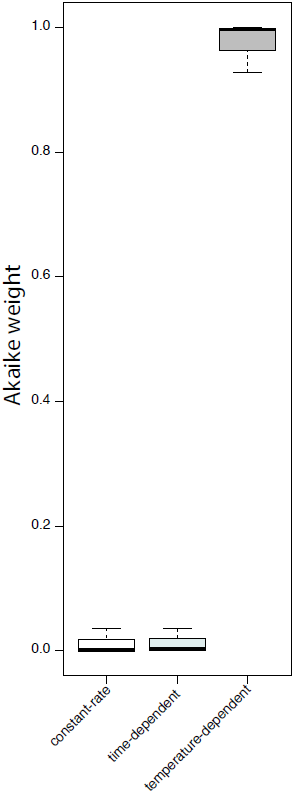
Akaike weights for constant-rate models, time-dependent models,and temperature-dependent models across all temperature-dependent (λ trees simulated witha high extinction rate ((λ = 0.75e^0.35t,^ (μ = 0.7).

**Supplemental Figure 5:**
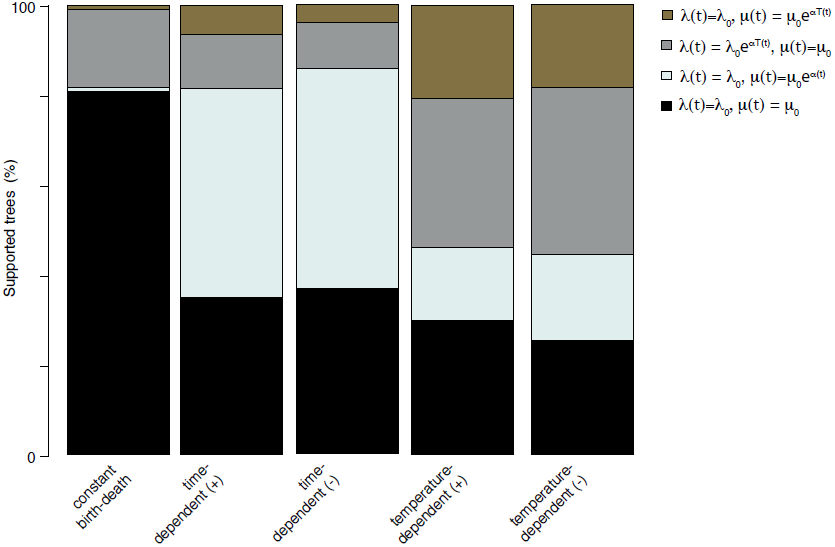
The ability to correctly recover trees simulated under constant-rate (λ = 0.2, (μ = 0.01), positive and negative dependence of (μ on time (λ_0_ = 0.5, (μ_0_ = 0.01, α_μ_ = ±0.4), and positive and negative dependence of (μ on temperature ((λ_0_ = 0.4, (μ_0_ = 0.1,α _μ_ = ±0.4). In addition to the simulating models, trees were fit with models with anexponential dependency of λ on temperature. Each column shows the percentage of trees simulated under the described models that find support.

**Supplemental Figure 6:**
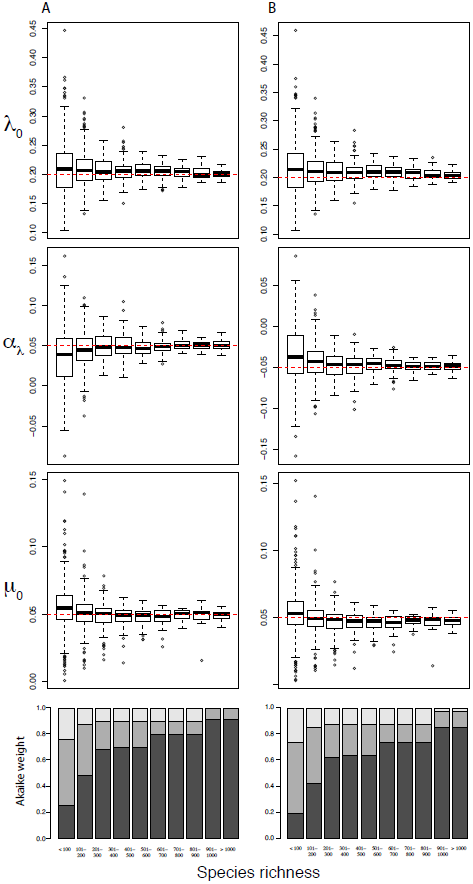
Parameter estimates for trees simulated with a positive (left) and negative (right) exponential dependency of λ on temperature by fitting the temperature-dependent model. Parameter estimates are shown for trees with different species richness. Simulated parameters are marked by dashed red lines. Akaike weights are shown for treesfitted with constant-rate models (light grey), time-dependent models (medium grey), and temperature-dependent models (dark grey). Weights are averaged across all trees within each species richness bracket.

**Supplemental Figure 7:**
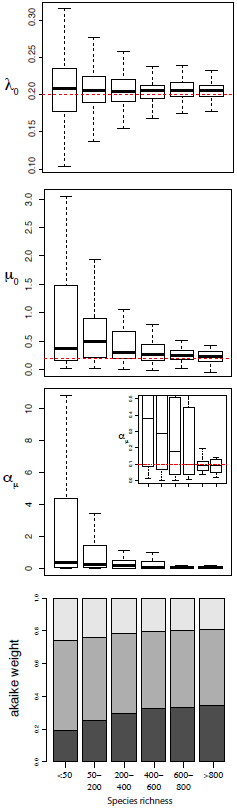
Parameter estimates for (A) λ_0_, (B) *μ0*, (C) αμ for trees simulated with a positive exponential dependency of *μ* on temperature by fitting the temperature-dependent model. Parameter estimates are shown for trees with different species richness. Simulated parameters are marked by dashed red lines. (C, inset) A magnified plot of estimates. (D) Akaike weights averaged over all trees for each species richness cohort for models fitted with a constant-rate model (light grey), time-dependent model (medium grey),and model with *μ* dependent on temperature (dark grey).

**Supplemental Figure 8:**
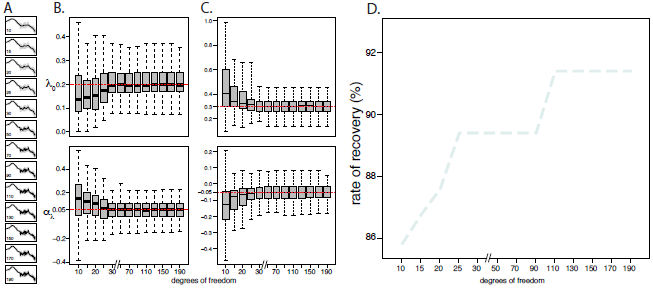
Model selection and parameter estimation in temperature-dependent trees under various spline smoothings in the fitted model. (A) Plots of temperature curve splines smoothed by different degrees of freedom. The time-series of temperature data are shown in grey dots and the smoothed curves in black lines. (B,C) Parameter estimates for trees simulated with an exponential dependency of speciation on temperature (B, λ = 0.2e^0.05T(t)^; C, λ = 0.3e^0.05T(t)^) with temperature curves determined using generalized cross-validation (degrees of freedom=208), where the fitted temperature-dependent models have temperature curve splines smoothed by different degrees of freedom. Simulated parameters are marked by dashed red lines. (D) The percentage of temperature-dependent trees, simulated with a temperature curve determined using generalized cross-validation, best supported by models fit with temperature curve splines smoothed by different degrees of freedom versus constant-rate models and time-dependent models with an exponential dependence on speciation.

**Supplemental Figure 9:**
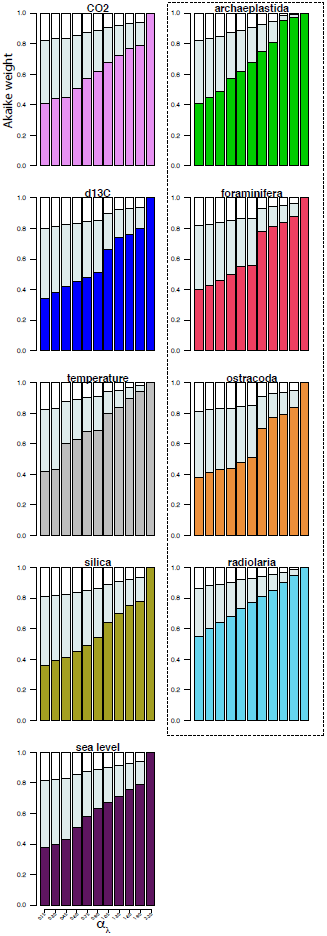
Mean of the Akaike weights for constant-rate models (light grey), time-dependent models (medium grey), and environment-dependent models (darkgrey) across all environment-dependent λ trees with the same α_λ_ (see Figure 3).

**Supplemental Figure 10.**
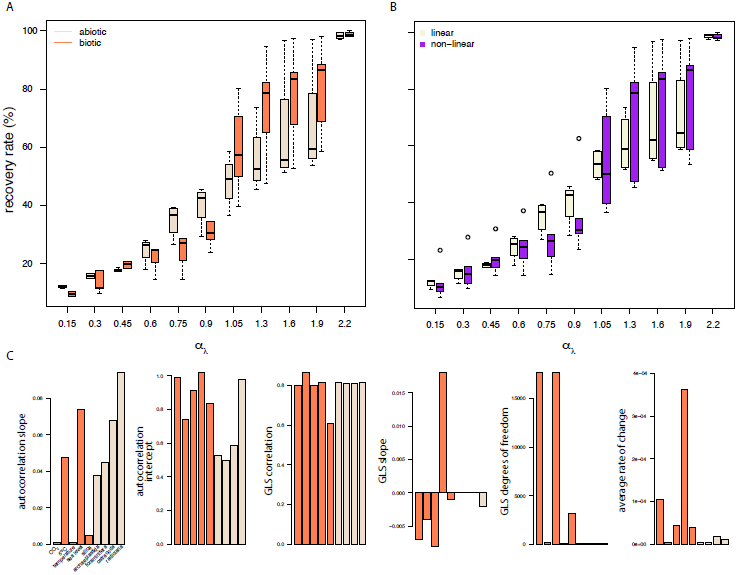
Effect of different characteristics on the rate of recovery of environmental curves. The ability to correctly recover the simulated model at differentvalues of α_λ_ for (A) abiotic and biotic variables (see Figure 3) and (B) linear and non-linear environmental curves. (C) Barplots for the slope and intercept for regression models fit to autocorrelation functions for multiple lag-times for each environmental curve; the correlation,slope, and degrees of freedom estimated for generalized least squares (GLS) linear fits to each environmental curve; and the average rate of change of each environmental variable with respect to time. Bars are colored according to panel A.

**Supplemental Figure 11:**
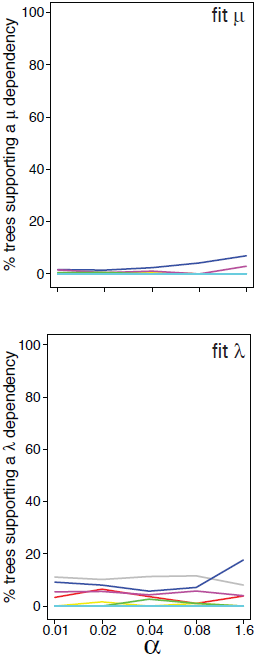
Rate of recovery for trees simulated with an exponential depen-dency of μ on different paleoenvironments, X, (μ= μ_0_e^αX(t)^) for varying values of α _μ_and constant λ. (Top) Simulated trees fitted with models with an exponential dependency ofμ on the paleoenvironment and a constant λ. (Bottom) Simulated trees fitted with models with an exponential dependency of λ on the paleoenvironment and a constant μ. Colors correspond to Figure 3.

**Supplemental Figure 12:**
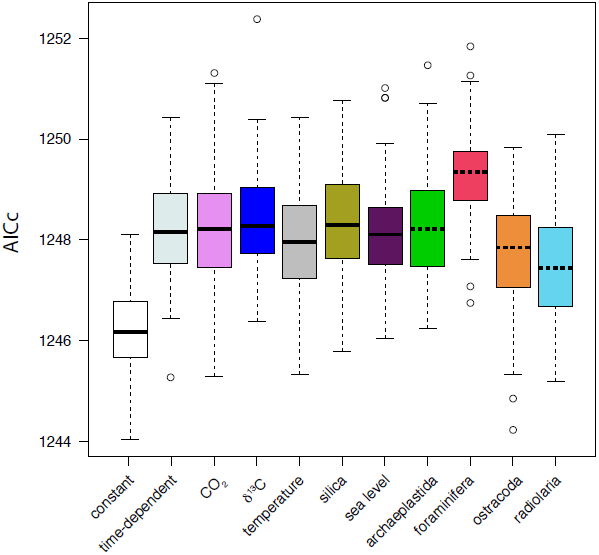
Environment-dependency in Ruminantia computed from the Ruminantia supertree (12). (A) AICc support for different environment-dependent models, a constant-rate birth-death model, and an exponential time-dependent model (without extinction) on a distribution of 5000 posteriorly sampled probabilities of the Ruminantia supertree. All environment-dependent models have an exponential dependency on the envi-ronmental variable. Colors and line type correspond to Figure 3.

## References

[1] J. Alroy. Geographical, environmental and intrinsic biotic controls on Phanerozoic marine diversification: Controls on phanerozoic marine diversification. Palaeontology, 53(6):1211–1235, Nov. 2010.

[2] P. M. B. Bacquet, O. Brattström, H.-L. Wang, C. E. Allen, C. Löfstedt, P. M. Brakefield, and C. M. Nieberding. Selection on male sex pheromone composition contributes to butterfly reproductive isolation. Proceedings. Biological Sciences, 282(1804):20142734, Apr. 2015.

[3] A. D. Barnosky. Distinguishing the effects of the red queen and court jester on miocene mammal evolution in the northern rocky mountains. Journal of Vertebrate Paleontology, 21(1):172–185, Mar. 2001.

[4] J. M. Beaulieu and B. C. O’Meara. Extinction can be estimated from moderately sized molecular phylogenies: BRIEF COMMUNICATION. Evolution, 69(4):1036–1043, Apr. 2015.

[5] D. J. Beerling and D. L. Royer. Convergent Cenozoic CO2 history. Nature Geoscience, 4(7):418–420, June 2011.

[6] C. J. P. Belisle. Convergence Theorems for a Class of Simulated Annealing Algorithms on d. Journal of Applied Probability, 29(4):885, Dec. 1992.

[7] M. Benton. Diversification and extinction in the history of life. Science, 268(5207):52, Apr. 1995.

[8] M. J. Benton. The Red Queen and the Court Jester: species diversity and the role of biotic and abiotic factors through time. Science (New York, N.Y.), 323(5915):728–732, Feb. 2009.

[9] F. Bibi.A multi-calibrated mitochondrial phylogeny of extant Bovidae (Artiodactyla, Ruminantia) and the importance of the fossil record to systematics. BMC Evolutionary Biology, 13(1):166, 2013.

[10] A. Buchan, G. R. LeCleir, C. A. Gulvik, and J. M. González. Master recyclers: features and functions of bacteria associated with phytoplankton blooms. Nature Reviews Microbiology, 12(10):686–698, Aug. 2014.

[11] K. P. Burnham and D. R. Anderson, editors. Model Selection and Multimodel Inference. Springer New York, New York, NY, 2004.

[12] J. L. Cantalapiedra, R. G. Fitzjohn, T. S. Kuhn, M. H. Fernández, D. DeMiguel, B. Azanza,J. Morales, and A. Ø. Mooers. Dietary innovations spurred the diversification of ruminants during the Caenozoic. Proceedings. Biological Sciences / The Royal Society, 281(1776):20132746, Feb. 2014.

[13] T. E. Cerling, J. M. Harris, B. J. MacFadden, M. G. Leakey, J. Quade, V. Eisenmann, and J. R. Ehleringer. Global vegetation change through the Miocene/Pliocene boundary. Nature, 389(6647):153–158, Sept. 1997.

[14] P. Cermeño, P. G. Falkowski, O. E. Romero, M. F. Schaller, and S. M. Vallina. Continental erosion and the Cenozoic rise of marine diatoms. Proceedings of the National Academy of Sciences, 112(14):4239–4244, Apr. 2015.

[15] Z.-Q. Chen and M. J. Benton. The timing and pattern of biotic recovery following the end-Permian mass extinction. Nature Geoscience, 5(6):375–383, May 2012.

[16] S. Claramunt and J. Cracraft. A new time tree reveals Earth history’s imprint on the evolution of modern birds. Science Advances, 1(11):e1501005, Dec. 2015.

[17] J. Clavel and H. Morlon. Accelerated body size evolution during cold climatic periods in the Cenozoic. Proceedings of the National Academy of Sciences of the United States of America, 114(16):4183–4188, Apr. 2017.

[18] C. Coiffard, B. Gomez, V. Daviero-Gomez, and D. L. Dilcher. Rise to dominance of angiosperm pioneers in European Cretaceous environments. Proceedings of the National Academy of Sciences of the United States of America, 109(51):20955–20959, Dec. 2012.

[19] F. L. Condamine, J. Rolland, and H. Morlon. Macroevolutionary perspectives to environmental change. Ecology Letters, pages 72–85, May 2013.

[20] F. L. Condamine, F. A. H. Sperling, N. Wahlberg, J.-Y. Rasplus, and G. J. Kergoat. What causes latitudinal gradients in species diversity? Evolutionary processes and ecological constraints on swallowtail biodiversity. Ecology Letters, 15(3):267–277, Mar. 2012.

[21] F. L. Condamine, E. F. A. Toussaint, A.-L. Clamens, G. Genson, F. A. H. Sperling, and G. J. Kergoat. Deciphering the evolution of birdwing butterflies 150 years after Alfred Russel Wallace. Scientific Reports, 5:11860, July 2015.

[22] J. Drury, J. Clavel, M. Manceau, and H. Morlon. Estimating the Effect of Competition on Trait Evolution Using Maximum Likelihood Inference. Systematic Biology, 65(4):700–710, July 2016.

[23] J. R. Ehleringer, T. Cerling, and M. D. Dearing. A history of atmospheric CO2 and its effects on plants, animals, and ecosystems. Springer Science & Business Media, 2006.

[24] P. R. Ehrlich and P. H. Raven. Butterflies and Plants: A Study in Coevolution. Evolution, 18(4):586, Dec. 1964.

[25] D. H. Erwin. Macroevolution of ecosystem engineering, niche construction and diversity. Trends in ecology & evolution, 23(6):304–310, June 2008.

[26] D. H. Erwin. Climate as a driver of evolutionary change. Current biology: CB, 19(14):R575–583, July 2009.

[27] R. S. Etienne, B. Haegeman, T. Stadler, T. Aze, P. N. Pearson, A. Purvis, and A. B. Phillimore. Diversity-dependence brings molecular phylogenies closer to agreement with the fossil record. Proceedings. Biological Sciences / The Royal Society, 279(1732):1300–1309, Apr. 2012.

[28] T. H. G. Ezard, T. Aze, P. N. Pearson, and A. Purvis. Interplay between changing climate and species’ ecology drives macroevolutionary dynamics. Science (New York, N.Y.), 332(6027):349–351, Apr. 2011.

[29] T. H. G. Ezard and A. Purvis. Environmental changes define ecological limits to species richness and reveal the mode of macroevolutionary competition. Ecology Letters, 19(8):899–906, Aug. 2016.

[30] T. H. G. Ezard, T. B. Quental, and M. J. Benton. The challenges to inferring the regulators of biodiversity in deep time. Philosophical Transactions of the Royal Society of London. Series B, Biological Sciences, 371(1691):20150216, Apr. 2016.

[31] R. G. FitzJohn. Diversitree: comparative phylogenetic analyses of diversification in R: *Diversitree*. Methods in Ecology and Evolution, 3(6):1084–1092, Dec. 2012.

[32] M. Foote, A. Miller, D. Raup, and S. Stanley. Principles of Paleontology. W. H. Freeman, 2007.

[33] L. M. Gerhart and J. K. Ward. Plant responses to low [CO2] of the past: Tansley review. New Phytologist, 188(3):674–695, Nov. 2010.

[34] G. H. Golub, M. Heath, and G. Wahba. Generalised cross-validation as a method for choosing a good ridge parameter. Technometrics, 21(2):215–223, 1979.

[35] B. Hannisdal and S. E. Peters. Phanerozoic Earth system evolution and marine biodiversity. Science (New York, N.Y.), 334(6059):1121–1124, Nov. 2011.

[36] B. Hansen. Testing for Linearity. Journal of Economic Surveys, 13(5):551–576, Dec. 1999.

[37] J. Hansen, M. Sato, G. Russell, and P. Kharecha. Climate sensitivity, sea level and atmospheric carbon dioxide. Philosophical Transactions of the Royal Society A: Mathematical, Physical and Engineering Sciences, 371(2001):20120294–20120294, Sept. 2013.

[38] S. Höhna. Fast simulation of reconstructed phylogenies under global time-dependent birth-death processes. Bioinformatics (Oxford, England), 29(11):1367–1374, June 2013.

[39] S. M. Holland and M. E. Patzkowsky. The stratigraphy of mass extinction. Palaeontology, 58(5):903–924, Sept. 2015.

[40] M. E. Katz, J. D. Wright, K. G. Miller, B. S. Cramer, K. Fennel, and P. G. Falkowski. Biological overprint of the geological carbon cycle. Marine Geology, 217(3):323–338, 2005.

[41] K. I. Keating. Blue-Green Algal Inhibition of Diatom Growth: Transition from Mesotrophic to Eutrophic Community Structure. Science, 199(4332):971–973, Mar. 1978.

[42] G. J. Kergoat, L. Soldati, A.-L. Clamens, H. Jourdan, R. Jabbour-Zahab, G. Genson, P. Bouchard, and F. L. Condamine. Higher level molecular phylogeny of darkling beetles (Coleoptera: Tenebrionidae): Darkling beetle phylogeny. Systematic Entomology, 39(3):486–499, July 2014.

[43] M. A. Knoll and W. C. James. Effect of the advent and diversification of vascular land plants on mineral weathering through geologic time. Geology, 15(12):1099–1102, Dec. 1987.

[44] D. Lazarus. Neptune: A marine micropaleontology database. Mathematical Geology, 26(7):817–832, Oct. 1994.

[45] D. Lazarus, J. Barron, J. Renaudie, P. Diver, and A. Türke. Cenozoic Planktonic Marine Diatom Diversity and Correlation to Climate Change. PLoS ONE, 9(1):e84857, Jan. 2014.

[46] L. H. Liow, T. Reitan, and P. G. Harnik. Ecological interactions on macroevolutionary time scales: clams and brachiopods are more than ships that pass in the night. Ecology Letters, 18(10):1030–1039, Oct. 2015.

[47] L. H. Liow, L. Van Valen, and N. C. Stenseth. Red Queen: from populations to taxa and communities. Trends in Ecology & Evolution, 26(7):349–358, July 2011.

[48] F. G. Marx and M. D. Uhen. Climate, Critters, and Cetaceans: Cenozoic Drivers of the Evolution of Modern Whales. Science, 327(5968):993–996, Feb. 2010.

[49] P. J. Mayhew, G. B. Jenkins, and T. G. Benton. A long-term association between global temperature and biodiversity, origination and extinction in the fossil record. Proceedings. Biological Sciences / The Royal Society, 275(1630):47–53, Jan. 2008.

[50] K. G. Miller. The Phanerozoic Record of Global Sea-Level Change. Science, 310(5752):1293–1298, Nov. 2005.

[51] G. G. Mittelbach, D. W. Schemske, H. V. Cornell, A. P. Allen, J. M. Brown, M. B. Bush, S. P. Harrison, A. H. Hurlbert, N. Knowlton, H. A. Lessios, C. M. McCain, A. R. McCune, L. A. McDade, M. A. McPeek, T. J. Near, T. D. Price, R. E. Ricklefs, K. Roy, D. F. Sax, D. Schluter, J. M. Sobel, and M. Turelli. Evolution and the latitudinal diversity gradient: speciation, extinction and biogeography. Ecology Letters, 10(4):315–331, Apr. 2007.

[52] H. Morlon. Phylogenetic approaches for studying diversification. Ecology letters, 17(4):508–525, Apr. 2014.

[53] H. Morlon, E. Lewitus, F. L. Condamine, M. Manceau, J. Clavel, and J. Drury. RPANDA: an R package for macroevolutionary analyses on phylogenetic trees. Methods in Ecology and Evolution, 7(5):589–597, May 2016.

[54] H. Morlon, T. L. Parsons, and J. B. Plotkin. Reconciling molecular phylogenies with the fossil record. Proceedings of the National Academy of Sciences of the United States of America, 108(39):16327–16332, Sept. 2011.

[55] S. Nee, E. C. Holmes, R. M. May, and P. H. Harvey. Extinction rates can be estimated from molecular phylogenies. Philosophical transactions of the Royal Society of London. Series B, Biological sciences, 344(1307):77–82, Apr. 1994.

[56] J. A. Nelder and R. Mead. A Simplex Method for Function Minimization. The Computer Journal, 7(4):308–313, Jan. 1965.

[57] S. L. Nuismer and L. J. Harmon. Predicting rates of interspecific interaction from phylogenetic trees. Ecology Letters, 18(1):17–27, Jan. 2015.

[58] G. Ocampo and J. T. Columbus. Molecular phylogenetics of suborder Cactineae (Caryophyllales), including insights into photosynthetic diversification and historical biogeography. American Journal of Botany, 97(11):1827–1847, Nov. 2010.

[59] A. M. Oehlert and P. K. Swart. Interpreting carbonate and organic carbon isotope covariance in the sedimentary record. Nature Communications, 5:4672, Aug. 2014.

[60] J.-Y. Park, J.-S. Kug, J. Bader, R. Rolph, and M. Kwon. Amplified Arctic warming by phytoplankton under greenhouse warming. Proceedings of the National Academy of Sciences, 112(19):5921–5926, May 2015.

[61] O. A. Pérez-Escobar, G. Chomicki, F. L. Condamine, A. P. Karremans, D. Bogarín, N. J. Matzke, D. Silvestro, and A. Antonelli. Recent origin and rapid speciation of neotropical orchids in the world’s richest plant biodiversity hotspot. New Phytologist, 215(2):891–905, 2017. 2017-23782.

[62] A. B. Phillimore and T. D. Price. Density-dependent cladogenesis in birds. PLoS biology, 6(3):e71, Mar. 2008.

[63] T. B. Quental and C. R. Marshall. Diversity dynamics: molecular phylogenies need the fossil record. Trends in Ecology & Evolution, 25(8):434–441, Aug. 2010.

[64] D. L. Rabosky. Extinction rates should not be estimated from molecular phylogenies. Evolution; international journal of organic evolution, 64(6):1816–1824, June 2010.

[65] D. L. Rabosky. Challenges in the estimation of extinction from molecular phylogenies: A response to Beaulieu and O’Meara: BRIEF COMMUNICATION. Evolution, 70(1):218–228, Jan. 2016.

[66] D. L. Rabosky and E. E. Goldberg. Model inadequacy and mistaken inferences of trait-dependent speciation. Systematic Biology, 64(2):340–355, Mar. 2015.

[67] D. M. Raup and J. J. Sepkoski. Mass extinctions in the marine fossil record. Science (New York, N.Y.), 215(4539):1501–1503, Mar. 1982.

[68] J. A. Raven and M. Giordano. Algae. Current Biology, 24(13):R590–R595, July 2014.

[69] J. M. Robinson. Speculations on carbon dioxide starvation, Late Tertiary evolution of stomatal regulation and floristic modernization. Plant, Cell and Environment, 17(4):345–354, Apr. 1994.

[70] K. Roy, G. Hunt, and D. Jablonski. Phylogenetic conservatism of extinctions in marine bivalves. Science (New York, N.Y.), 325(5941):733–737, Aug. 2009.

[71] N. Seddon, C. A. Botero, J. A. Tobias, P. O. Dunn, H. E. A. Macgregor, D. R. Rubenstein, J. A. C. Uy, J. T. Weir, L. A. Whittingham, and R. J. Safran. Sexual selection accelerates signal evolution during speciation in birds. Proceedings. Biological Sciences, 280(1766):20131065, Sept. 2013.

[72] J. J. Sepkoski. A kinetic model of Phanerozoic taxonomic diversity; III, Post-Paleozoic families and mass extinctions. Paleobiology, 10(2):246–267, 1984.

[73] D. Silvestro, J. Schnitzler, L. H. Liow, A. Antonelli, and N. Salamin. Bayesian estimation of speciation and extinction from incomplete fossil occurrence data. Systematic Biology, 63(3):349–367, May 2014.

[74] M. Slatkin. Gene flow and the geographic structure of natural populations. Science (New York, N.Y.), 236(4803):787–792, May 1987.

[75] A. B. Smith. Marine diversity through the Phanerozoic: problems and prospects. Journal of the Geological Society, 164(4):731–745, July 2007.

[76] J. M. Smith. The Causes of Extinction. Philosophical Transactions of the Royal Society of London. B, Biological Sciences, 325(1228):241, Nov. 1989.

[77] R. A. Spicer and J. L. Chapman. Climate change and the evolution of high-latitude terrestrial vegetation and floras. Trends in Ecology & Evolution, 5(9):279–284, Sept. 1990.

[78] M. E. Steeman, M. B. Hebsgaard, R. E. Fordyce, S. Y. W. Ho, D. L. Rabosky, R. Nielsen, C. Rahbek, H. Glenner, M. V. Sorensen, and E. Willerslev. Radiation of Extant Cetaceans Driven by Restructuring of the Oceans. Systematic Biology, 58(6):573–585, Dec. 2009.

[79] Y. Teslenko. Some aspects of evolution of terrestrial plants. Geol. Geofizica (Novosibirsk), 11:58–64, 1967.

[80] L. Van Valen. A new evolutionary law. Evolutionary Theory, 1:1–30, 1973.

[81] K. L. Voje, Ø. H. Holen, L. H. Liow, and N. C. Stenseth. The role of biotic forces in driving macroevolution: beyond the Red Queen. Proceedings of the Royal Society B: Biological Sciences, 282(1808), May 2015.

[82] T. Volk. Rise of angiosperms as a factor in long-term climatic cooling. Geology, 17(2):107–110, Feb. 1989.

[83] A. von Humboldt. Ansichten der Natur: mit wissenschaftliehen Erl¨auterungen, volume 1. JG Cotta, 1860.

[84] E. S. Vrba. Turnover-pulses, the Red Queen, and related topics. American Journal of Science, 293(A):418–452, Jan. 1993.

[85] A. R. Wallace. Tropical nature, and other essays. Macmillan and Company, 1878.

[86] H. Whitehead, B. McGill, and B. Worm. Diversity of deep-water cetaceans in relation to temperature: implications for ocean warming. Ecology Letters, Sept. 2008.

[87] A. M. Winkler, P. Kochunov, J. Blangero, L. Almasy, K. Zilles, P. T. Fox, R. Duggirala, and D. C. Glahn. Cortical thickness or grey matter volume? The importance of selecting the phenotype for imaging genetics studies. NeuroImage, 53(3):1135–1146, Nov. 2010.

[88] S. Wright. Evolution in Mendelian Populations. Genetics, 16(2):97–159, Mar. 1931.

[89] J. C. Zachos, G. R. Dickens, and R. E. Zeebe. An early Cenozoic perspective on greenhouse warming and carbon-cycle dynamics. Nature, 451(7176):279–283, Jan. 2008.

